# PRMT5 epigenetically regulates the E3 ubiquitin ligase ITCH to influence lipid accumulation during mycobacterial infection

**DOI:** 10.1101/2021.11.09.467864

**Authors:** Salik Miskat Borbora, R.S. Rajmani, Kithiganahalli Narayanaswamy Balaji

**Affiliations:** Department of Microbiology and Cell Biology, Indian Institute of Science, Bangalore - 560012, Karnataka, India; Center for Infectious Disease Research, Indian Institute of Science, Bangalore – 560012, Karnataka, India

## Abstract

*Mycobacterium tuberculosis* (Mtb), the causative agent of tuberculosis (TB), triggers enhanced accumulation of lipids to generate foamy macrophages (FMs). This process has been often attributed to the surge in the expression of lipid influx genes with a concomitant decrease in those involved in lipid efflux genes. Here, we define an Mtb-orchestrated modulation of the ubiquitination mechanism of lipid accumulation markers to enhance lipid accretion during infection. We find that Mtb infection represses the expression of the E3 ubiquitin ligase, ITCH, resulting in the sustenance of key lipid accrual molecules viz. ADRP and CD36, that are otherwise targeted by ITCH for proteasomal degradation. In line, overexpressing ITCH in Mtb-infected cells was found to suppress Mtb-induced lipid accumulation. Molecular analyses including loss-of-function and ChIP assays demonstrated a role for the concerted action of the transcription factor YY1 and the arginine methyl transferase PRMT5 in restricting the expression of *Itch* gene by conferring repressive symmetrical H4R3me2 marks on its promoter. Consequently, siRNA-mediated depletion of YY1 or PRMT5 rescued ITCH expression, thereby compromising the levels of Mtb-induced ADRP and CD36 and limiting FM formation during infection. Accumulation of lipids within the host has been implicated as a pro-mycobacterial process that aids in pathogen persistence and dormancy. In our study, perturbation of PRMT5 enzyme activity resulted in compromised lipid levels and reduced mycobacterial survival in primary murine macrophages (*ex vivo*) and in a therapeutic mouse model of TB infection (*in vivo*). These findings provide new insights on the role of PRMT5 and YY1 in augmenting mycobacterial pathogenesis. Thus, we posit that our observations could help design novel adjunct therapies and combinatorial drug regimen for effective anti-TB strategies.

**Author Summary:** *Mycobacterium tuberculosis* generates lipid-laden cells (foamy macrophages-FMs) that offer a favorable shelter for its persistence. During infection, we observe a significant reduction in the expression of the E3 ubiquitin ligase, ITCH. This repression allows the sustenance of key lipid accretion molecules (ADRP and CD36), by curbing their proteasomal degradation. Further, we show the repression of ITCH to be dependent on the concerted action of the bifunctional transcription factor, YY1 and the arginine methyl transferase, PRMT5. NOTCH signaling pathway was identified as a master-regulator of YY1 expression. *In vitro* and *in vivo* analyses revealed the significance of PRMT5 in regulating FM formation and consequently mycobacterial burden.

## Introduction

*Mycobacterium tuberculosis* (Mtb), the principal etiological agent of the pulmonary ailment, tuberculosis (TB), continues to co-evolve with the human population making itself one of the major infectious diseases afflicting mankind. Globally in 2020, there were an estimated 1.5 million deaths due to TB, a figure which saw a sharp increment following the onset of the COVID-19 pandemic (WHO, Annual TB Report, 2021).

Upon infecting the host, Mtb directs the release of definite cues that aids in the formation of a highly organized immune structure-the granuloma (1). TB granulomas are constituted by macrophages, neutrophils, monocytes, dendritic cells, B- and T-cells, fibroblasts, and epithelial cells (2, 3). Anatomical dissection into the distinct features of the macrophages within the TB granuloma has revealed accumulated lipids as one of the leading physiological countenances. Also referred to as foamy macrophages (FM), these cells act as a protective niche wherein the bacterium thrives and persists. Mtb deregulates host cellular machinery to trigger increased lipid intake into the cells with a concomitant decrease in the efflux of lipids-thereby contributing to a lipid-rich environment (4). While the transcriptional regulation of a gamut of such molecules has been reported, there are scant reports on the post-translational regulation of proteins necessary for FM formation in the context of Mtb infection. Parallel studies have highlighted deregulated autophagy (5) and lipophagy (6) to be instrumental in regulating lipid turnover during Mtb infection. Ubiquitin-dependent degradation of specific proteins implicated during FM formation is yet another mechanism that can control the stability of proteins that aid in lipid accumulation, thereby securing the integrity of lipids (7). In this context, Nedd4-family ligases (a sub-family of HECT-domain containing E3 ligases), among others, are known to play distinctive roles in maintaining the stability of diverse proteins involved in FM formation. Notably, NEDD4-1 can interact with the ATP-Binding Cassette (ABC) transporters-ABCG1 and ABCG4, two essential regulators of cellular lipid homeostasis (8). Besides, another report alluded to the role of AIP4 or ITCH in maintaining the levels of the important FM protein, ADRP (9).

During TB, accumulated host lipids have been associated with mycobacterial survival as they provide essential nutrients, contribute to reduced antigen presentation (10–12) and aid in pathogen latency and reactivation (13). While it was our interest to define ubiquitination mechanisms contributing to copious levels of lipid within the cells, a study revealed that knocking down *Nedd4* in THP-1 macrophages elicited higher CFU of Mtb and other intracellular pathogens (14). This encouraged us to investigate the role of NEDD4-family of E3 ligases in the context of Mtb-driven FM formation and thereupon pathogen survival.

Pathogen-driven epigenetic changes have been indicated to govern the expression of distinct molecules that contribute to mycobacterial survival (4, 15–18). Molecular dissection of the specific roles of such epigenetic regulators bears significance as perturbing the same could provide valuable cues to modify existing drug regimen against TB. Emerging literature have indicated towards epigenetic regulation of NEDD4-family ligases. Precisely, HDAC2-dependent downregulation of histone methyltransferase Ehmt2 (G9a) contributed to the activation of NEDD4 due to the loss of repressive methylation signature (19). Thus, investigation on the contribution of epigenetics towards the expression of E3 ligases and consequently, FM generation during mycobacterial infection, seemed an intriguing question.

While a plethora of epigenetic regulators have been associated with the progression of several infectious diseases, including TB (4, 16, 18, 20, 21), there is a dearth of reports on the role of arginine methyl transferases such as Protein arginine methyl transferases (PRMTs) during distinct infections. Of all the characterized PRMTs, PRMT1, PRMT2, PRMT3, PRMT6, PRMT8 and CARM1 catalyze asymmetric methylation, while PRMT5 and PRMT9 bring about symmetric methylation on specific arginine (22). Available reports have demonstrated the role of PRMT5 in regulating lipid biogenesis. Notably, PRMT5 arbitrates seipin-mediated lipid droplet biogenesis in adipocytes (23). Besides, PRMT5 could enhance lipogenesis in cancer cells by methylating SREBP1a (24). These reports encouraged us to investigate a PRMT5-mediated regulation of FM formation during Mtb infection. Also, using specific inhibitors against PRMT5, we attempted to comprehend the possible roles of lipid dysregulation in augmenting mycobacterial survival *in vitro* and *in vivo.*

Here, we demonstrate that Mtb infection contributed to the repression of the E3 ligase, ITCH, in a NOTCH signaling-dependent manner, that leads to enhanced lipid accumulation in the macrophages. We observed that diminished levels of ITCH aided in sustaining the levels of key LD molecules, ADRP and CD36, and consequently elevating lipid pools within the cells. Additionally, we report the critical role of PRMT5 in mediating ITCH repression during virulent Mtb infection whilst underscoring the beneficial impact of the use of PRMT5 inhibitor in resolving tuberculous granuloma-like lesions in mice. Thus, the current study emphasizes on the crucial role of host epigenetic regulators in augmenting TB pathology and draws attention towards specific cellular nodes that could be targeted for the development of host-directed therapeutics against TB.

## Results

### Mycobacterial infection elicits differential expression of the E3 ubiquitin ligase, ITCH, to enhance lipid accumulation in cells

As distinct bona fide cellular organelles, lipid droplets are formed by the coordinated action of diverse processes which include its biogenesis, maturation, and turnover (25). As introduced, ubiquitination has been implicated in the regulation of proteins that contribute to lipid turnover within cells (26), thereby providing distinct regulatory potential to the entire process of lipid accumulation (27). Besides, Nedd4-family ligases were reported to be responsible for ubiquitination and degradation of lipid-associated proteins such as ADRP (9). With this premise, we screened for Nedd4-family ligases that are differentially expressed during Mtb infection and can potentially aid in FM formation during mycobacterial infection. Interestingly, we found the transcript levels of the E3 ligase *Itch* to be significantly downregulated both *in vitro* **(Fig. S1A)** and *in vivo* **(Fig. S1B)**, prompting us to investigate the molecule further. This was corroborated at the protein level as primary murine macrophages displayed diminished levels of ITCH protein **(Fig. 1A)** upon mycobacterial infection, while homogenates from the lungs of Mtb-infected mice exhibited a similar downregulated expression of ITCH **(Fig. 1B)**.

**Figure 1.**
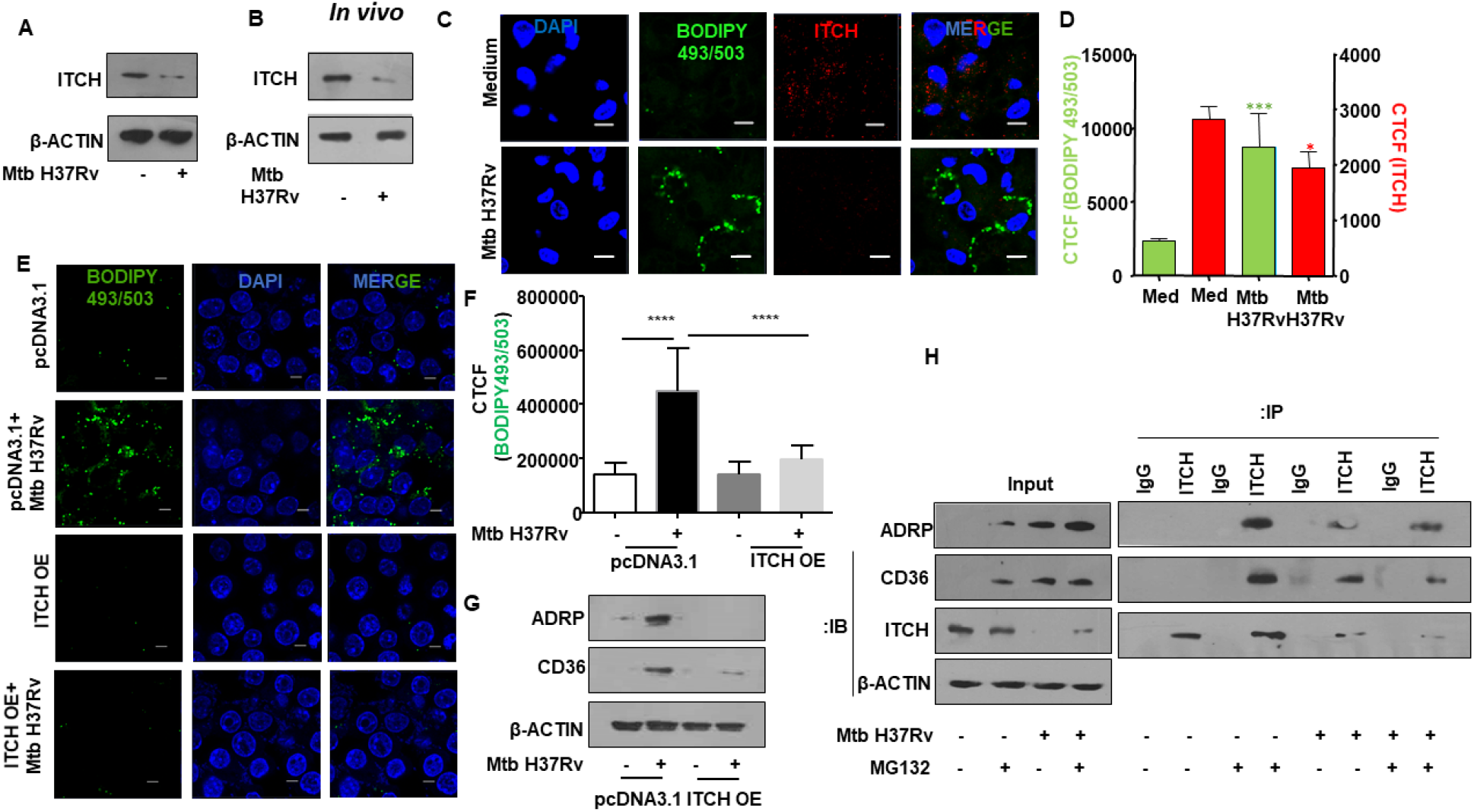
E3 ubiquitin ligase ITCH (AIP4) repression aids in Mtb-mediated enhanced lipid accumulation. **(A)** Mouse peritoneal macrophages were infected with Mtb H37Rv for 24 h. Whole cell lysates were assessed for ITCH protein expression by Western blotting. **(B)** Balb/c mice were aerosol-infected with Mtb H37Rv for 28 days. Protein levels of ITCH was analyzed in the lung homogenates of uninfected and infected mice by western blotting. **(C, D)** Mouse peritoneal macrophages were infected with Mtb H37Rv for 48 h and assessed for lipid accumulation (BODIPY493/503) and ITCH expression by immunofluorescence microscopy; **(C)** representative image and **(D)** respective quantification. **(E, F)** RAW 264.7 cells were transfected with pcDNA3.1 and ITCH OE vectors. Transfected cells were infected with Mtb H37Rv for 48 h and assessed for the accumulation of lipids by fluorescence microscopy (BODIPY493/503); **(E)** representative image and **(F)** respective quantification. **(G)** RAW 264.7 cells were transfected with pcDNA3.1 and ITCH OE vectors; Transfected cells were infected with Mtb H37Rv for 24 h and whole cell lysates were assessed for the levels of the indicated lipid accumulation markers by immunoblotting. **(H)** Mouse peritoneal macrophages were infected with Mtb H37Rv for 24 h and treated with MG132 for the last 4 h of infection. Whole cell lysates were immunoprecipitated with IgG control or ITCH antibodies and assessed for their interaction with the indicated lipid accumulation markers by immunoblotting. All immunoblotting and immunofluorescence data are representative of three independent experiments. Lung homogenates from at least three mice were independently assessed for ITCH expression by immunoblotting. Immunoblotting data is representative of three independent experiments. β-ACTIN was utilized as loading control. Med, medium; OE, over expression; CTCF, corrected total cell fluorescence. *, p<0.05; *** p<0.001; ****, p < 0.0001 (Student’s t-test in D, One way ANOVA in F; GraphPad Prism 6.0). Scale bar, 5 μm.

An earlier report had highlighted the role of ITCH in ubiquitinating ADRP, a key molecule during FM formation (9). Additionally, previous investigation had underscored the role of ADRP and CD36 in Mtb-mediated FM formation (4). To evaluate the role of ITCH in FM formation during Mtb infection, murine peritoneal macrophages (Mtb-infected and uninfected) were co-stained for lipid droplets and ITCH. Mtb infection revealed enhanced accumulation of lipids with a concomitant decrease in the levels of ITCH expression **(Fig. 1 C, D)**. Also, transient over expression of ITCH in RAW 264.7 macrophages compromised the Mtb-induced FM formation, thereby emphasizing on the role of ITCH in lipid accumulation **(Fig. 1E, F)**. Furthermore, ITCH over expression compromised the elevated levels of key lipid accumulation molecules, ADRP and CD36 **(Fig. 1G)**. These observations led us to assess the possible role of ITCH in the proteasomal degradation of the said proteins. We found a significant degree of interaction of ITCH with ADRP and CD36 in the uninfected scenario **(Fig. 1H)**, suggesting the contribution of ITCH in maintaining lipid homeostasis in macrophages under basal conditions. However, in the event of ITCH repression during Mtb infection, ADRP and CD36 levels are sustained, thereby leading to lipid accumulation. Together, these observations indicate a role for ITCH in regulating FM formation during Mtb infection.

### Transcription regulator, YY1, aids in Mtb-mediated repression of ITCH

Having established the repression of ITCH upon mycobacterial infection, we analyzed the promoter region of *Itch* gene to identify the potential regulators of ITCH expression. Bioinformatic assessment of the 2kb upstream sequence revealed two putative binding sites of the transcriptional regulator, YY1. A previous report suggests an enhanced expression of YY1 in human TB samples, where it plays a cardinal role in the regulation of specific cytokines (28). Essentially, YY1 is a zinc finger domain containing protein that can activate or repress genes based on its interacting partners (29). Primary murine macrophages displayed an enhanced expression of YY1 in a time-dependent manner upon mycobacterial infection **(Fig. 2A)**. A similar trend was observed *in vivo* as an elevated YY1 expression was observed in the lung homogenates from the lungs of Mtb-infected mice **(Fig. 2B)**. Analysis of the subcellular localization of YY1 in murine macrophages revealed that Mtb infection enhanced the nuclear localization of YY1 **(Fig. 2C)**, underscoring the possible role of YY1 in mediating transcriptional processes upon Mtb infection. Next, we compromised *Yy1* levels in murine macrophages with targeted siRNA and found a marked rescue in the Mtb-driven repression of ITCH **(Fig. 2D)**. This was further confirmed by ChIP assay, that indicated an enhanced recruitment of YY1 at its respective binding sites on the promoter of ITCH **(Fig. 2E)**.

**Figure 2.**
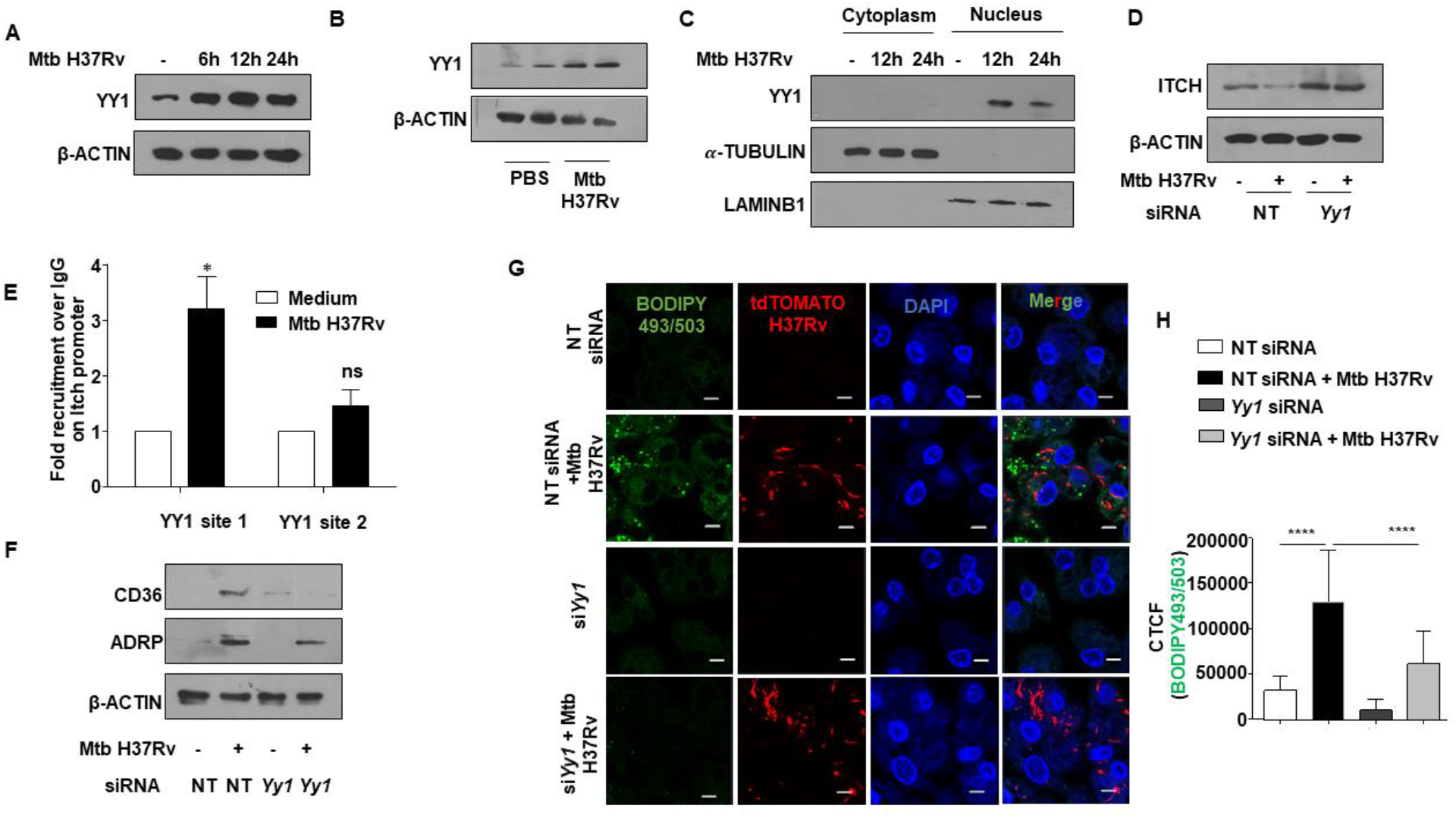
Transcription factor YY1 regulates ITCH repression during Mtb infection. **(A)** Mouse peritoneal macrophages were infected with Mtb H37Rv for the indicated time points and YY1 expression was analyzed in the whole cell lysates by immunoblotting. **(B)** Mouse peritoneal macrophages were infected with Mtb H37Rv for the indicated time points. Cells were partitioned into nuclear and cytoplasmic fractions and assessed for the nuclear translocation of YY1 by immunoblotting. **(C)** Balb/c mice were aerosol-infected with Mtb H37Rv for 28 days. Protein levels of YY1 was analyzed in the lung homogenates of uninfected and infected mice by western blotting. **(D)** Mouse peritoneal macrophages were transfected with NT or *Yy1* siRNAs. Transfected cells were infected with Mtb H37Rv for 24 h and assessed for the expression of ITCH by immunoblotting. **(E)** Mouse peritoneal macrophages were infected with Mtb H37Rv for 24 h and assessed for the recruitment of YY1 over the *Itch* promoter by ChIP assay. **(F)** Mouse peritoneal macrophages were transfected with NT or *Yy1* siRNAs. Transfected cells were infected with Mtb H37Rv for 24 h and assessed for the expression of ADRP and CD36 by immunoblotting. **(G, H)** Mouse peritoneal macrophages were transfected with NT or *Yy1* siRNAs. Transfected cells were infected with tdTomato Mtb H37Rv for 48 h and analyzed for lipid accumulation (BODIPY493/503) by confocal microscopy; **(G)** representative image and **(H)** respective quantification. All immunoblotting, immunofluorescence and ChIP data are representative of three independent experiments. Lung homogenates from at least three mice were independently assessed for YY1 expression by immunoblotting. β-ACTIN, *α*-TUBULIN and LAMINB1 were utilized as loading control. NT, non-targeting; CTCF, corrected total cell fluorescence; ChIP, Chormatin immunoprecipitation; ns, non-significant. *, p<0.05; ****, p < 0.0001 (Student’s t-test in E, One way ANOVA in H; GraphPad Prism 6.0). Scale bar, 5 μm.

With the premise of YY1-dependent downregulation of ITCH, we verified the contribution of YY1 in FM generation during Mtb infection. We found the expression of the FM markers, ADRP and CD36, to be compromised in Mtb-infected primary murine macrophages transfected with *Yy1* siRNA **(Fig. 2F)**. Furthermore, depletion of *Yy1* in primary murine macrophages also reduced lipid accumulation in Mtb-infected macrophages as assessed by the abundance of the BODIPY 493/503 signal **(Fig. 2G, H)**.

### NOTCH signaling pathway contributes to mycobacteria-induced lipid accumulation through YY1

Orchestrated cellular events such as FM formation are dictated by definite signals that might be either intracellular or extracellular. Studies have revealed that TNF receptor signaling through the downstream activation of the caspase cascade and the mammalian target of rapamycin complex 1 (mTORC1) is central to triglyceride accumulation in human macrophages infected with Mtb (30). Besides, NOTCH signaling and EGFR signaling have been separately implicated in Mtb-driven LD formation (4, 6).

Here, we sought to examine such upstream regulatory events that can potentially contribute to the differential expression of YY1 during mycobacterial infection. Using specific pharmacological interventions against the said pathways, we found the possible role of NOTCH signaling in the expression of YY1 as inhibition of the pathway using γ-secretase inhibitor (GSI) compromised the ability of Mtb to induce the expression of YY1 in macrophages **(Fig. 3A)**. NOTCH pathway activation is characterized by the cleavage of the intracellular domain of the NOTCH receptor (NICD), that shuttles to the nucleus to generate specific transcriptional response. In line with previous reports (31, 32), we found an activation of NOTCH signaling pathway (elevated NICD expression) in primary murine macrophages infected with Mtb **(Fig. S3A)**. The transcription factor, HES1, a bona fide target of NOTCH signaling, was found to be upregulated upon Mtb infection as assessed by transcript analysis **(Fig. S3B)** as well as luciferase reporter assay **(Fig. S3C)**.

**Figure 3.**
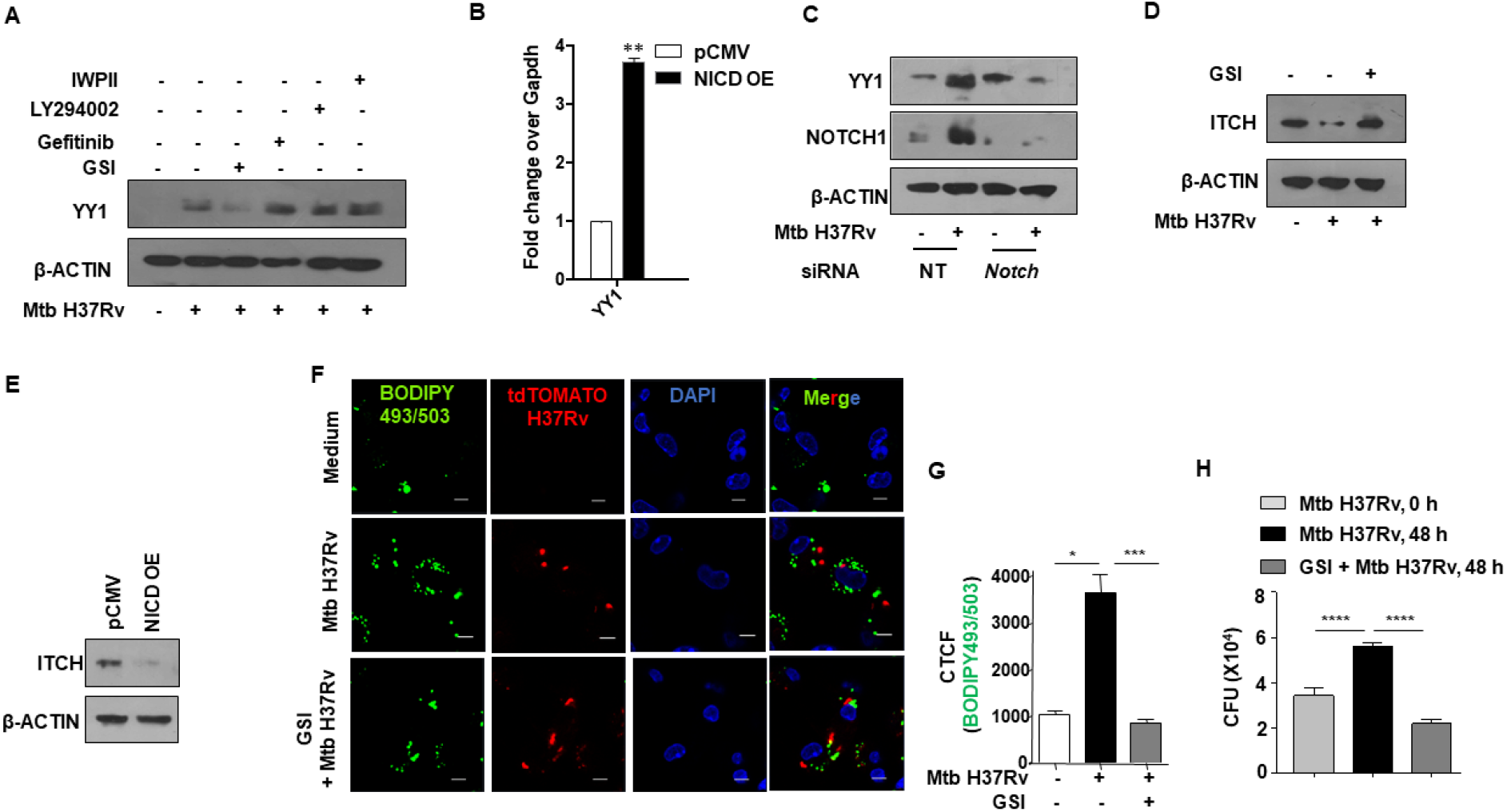
Activated NOTCH signaling pathway aids in enhanced YY1 production during Mtb infection. **(A)** Mouse peritoneal macrophages were treated with the indicated inhibitors for 1 h, followed by 24 h infection with Mtb H37Rv. Whole cell lysates were assessed for the expression of YY1 by immunoblotting. **(B)** RAW 264.7 cells were transfected with pCMV or NICD overexpression vectors and assessed for the transcript levels of *Yy1* by qRT-PCR. **(C)** Mouse peritoneal macrophages were transfected with NT or *Notch* siRNAs. Transfected cells were infected with Mtb H37Rv for 24 h and assessed for the expression of YY1 and NOTCH1 by immunoblotting. **(D)** Mouse peritoneal macrophages were treated with NOTCH pathway inhibitor GSI for 1 h, followed by 24 h infection with Mtb H37Rv. Whole cell lysates were assessed for the expression of ITCH by immunoblotting. **(E)** RAW 264.7 cells were transfected with control or NICD overexpression vectors and assessed for the expression of ITCH by immunoblotting. **(F, G)** Mouse peritoneal macrophages were treated with the NOTCH pathway inhibitor GSI for 1 h, followed by 48 h infection with tdTomato Mtb H37Rv and analyzed for lipid accumulation (BODIPY493/503) by confocal microscopy; **(F)** representative image and **(G)** respective quantification. **(H)** Mouse peritoneal macrophages were infected with Mtb H37Rv for 4 h. Extracellular bacteria were removed, and the infected cells were cultured in the presence or absence of PRMT5 inhibitor, EPZ015666 for 48 h. Cells were lysed and plated on 7H11 to enumerate intracellular Mtb H37Rv burden. qRT-PCR data represents mean±S.E.M. and immunoblotting and immunofluorescence data are representative of three independent experiments. OE, over expression; NT, non-targeting; CTCF, corrected total cell fluorescence; CFU, colony forming units. IWPII, WNT pathway inhibitor; LY294002, PI3K pathway inhibitor; Gefitinib, EGFR pathway inhibitor; GSI, Gamma secretase inhibitor (GSI), NOTCH pathway inhibitor. *, p<0.05; **, p<0.01; *** p<0.001; ****, p < 0.0001 (Student’s t-test in D, One way ANOVA in G,H; GraphPad Prism 6.0). Scale bar, 5 μm.

To verify the role of NOTCH pathway activation in augmenting YY1 levels, we overexpressed NICD in RAW 264.7 macrophages and found enhanced levels of YY1 transcript even in the absence of Mtb infection, underscoring the role of NOTCH in the elevated expression of YY1 **(Fig. 3B)**. Further, Mtb-mediated expression of YY1 was compromised in macrophages expressing *Notch1* siRNA **(Fig. 3C)**. Additionally, perturbation of the NOTCH pathway in murine peritoneal macrophages using GSI alleviated the Mtb-mediated diminished expression of ITCH **(Fig. 3D)**. Also, NICD overexpressing macrophages displayed compromised expression of ITCH **(Fig. 3E)**, corroborating our observations on the role of the NOTCH pathway in regulating the levels of YY1 and consequently the E3 ligase, ITCH. Furthermore, inhibition of the NOTCH pathway hindered the accumulation of lipids during Mtb infection **(Fig. 3F, G)**, thereby endorsing the significant role of NOTCH-YY1-ITCH axis in Mtb-induced FM formation.

The accumulation of host lipids during mycobacterial infection has been majorly attributed to the facilitation of nutrients to the bacteria. In this context, using *in vitro* CFU analysis, we found that perturbation of NOTCH pathway (using GSI) severely compromised mycobacterial survival in primary murine macrophages by rescuing the levels of ITCH and initiating ITCH-mediated ubiquitination of FM markers **(Fig. 3H)**. It has been already reported that NOTCH signaling aids in mycobacterial survival within the host (31).

### PRMT5 imparts repressive methylation signature on the promoter of ITCH during mycobacterial infection

As introduced, YY1 is a bifunctional transcription factor that is capable of activating or repressing genes based on its association with distinct factors. In the given scenario, since the expression of ITCH was repressed, we sought to delineate the possible way in which YY1 could accomplish the same. Ample evidence has highlighted the association of YY1 with the Polycomb-group (PcG) proteins to bring about repression of target genes. Notably, upon binding to the DNA, YY1 could initiate PcG protein recruitment that results in concomitant histone deacetylation and methylation (32). Herein, hinging on the available reports, we surmised if YY1 could bring about repression of ITCH by recruiting members of the PcG proteins. A ChIP assay was performed to assess for the recruitment of the principal methyl transferase, EZH2-a member of the PcG proteins, at the YY1 binding site on the promoter of ITCH. It was observed that Mtb infection did not elicit an appreciable recruitment of EZH2 or its cognate methylation signature H3K27me3 at the YY1 binding sites on the promoter of ITCH **(Fig. 4A)**. Based on this result, we construed that YY1 might interact with distinct epigenetic molecules other than EZH2 to bring about the repression of ITCH. Extensive review of literature revealed that PRMT1, an arginine methyl transferase could interact with YY1 to elicit the enhanced expression of genes (33). This hinted at a possible interaction between YY1 and enzyme(s) from the PRMT-family of methyl transferases. It is important to note that PRMT1 effectuates asymmetric dimethylation in the H4R3 residue (H4R3me2a) to initiate the expression of genes. Interestingly, symmetric dimethylation at the same arginine residue of histone 4 (H4R3me2s) brings about repression in gene expression. This symmetric dimethylation is catalyzed by the enzyme PRMT5 (34). This coerced us to conjecture if YY1 could interact with PRMT5 and catalyze the symmetric dimethylation (H4R3me2s) signature on the promoter region of ITCH and arbitrate its repression. A ChIP assay confirmed the occupancy of PRMT5 and its cognate repressive methylation mark at the YY1-binding sites on the promoter of ITCH **(Fig. 4B)**. We also observed that upon the perturbation of PRMT5 enzymatic activity with a specific inhibitor EPZ015666, Mtb-mediated diminished expression of ITCH was rescued in primary murine macrophages **(Fig. 4C)**. Besides, gene-specific knockdown of *Prmt5* in primary murine macrophages alleviated Mtb-mediated repression of ITCH during infection **(Fig. 4D)**. Further, immunoprecipitation analysis in murine macrophages revealed enhanced interaction between YY1 and PRMT5 upon mycobacterial infection **(Fig. 4E)**, implying a possible scenario wherein YY1 could engage PRMT5 on the ITCH promoter, resulting in the deposition of repressive methylation mark H4R3me2s. To conclusively implicate the significance of the association of YY1 and PRMT5 for the recruitment of the repressive methylation signature on the promoter of ITCH, siRNA against *Yy1* was employed to selectively decrease the levels of *Yy1* in primary murine macrophages. Subsequently, it was observed that cells which were depleted of *Yy1* showed compromised recruitment of PRMT5 as well as the repressive methylation mark H4R3me2s on the promoter of ITCH **(Fig. 4F)**.

**Figure 4.**
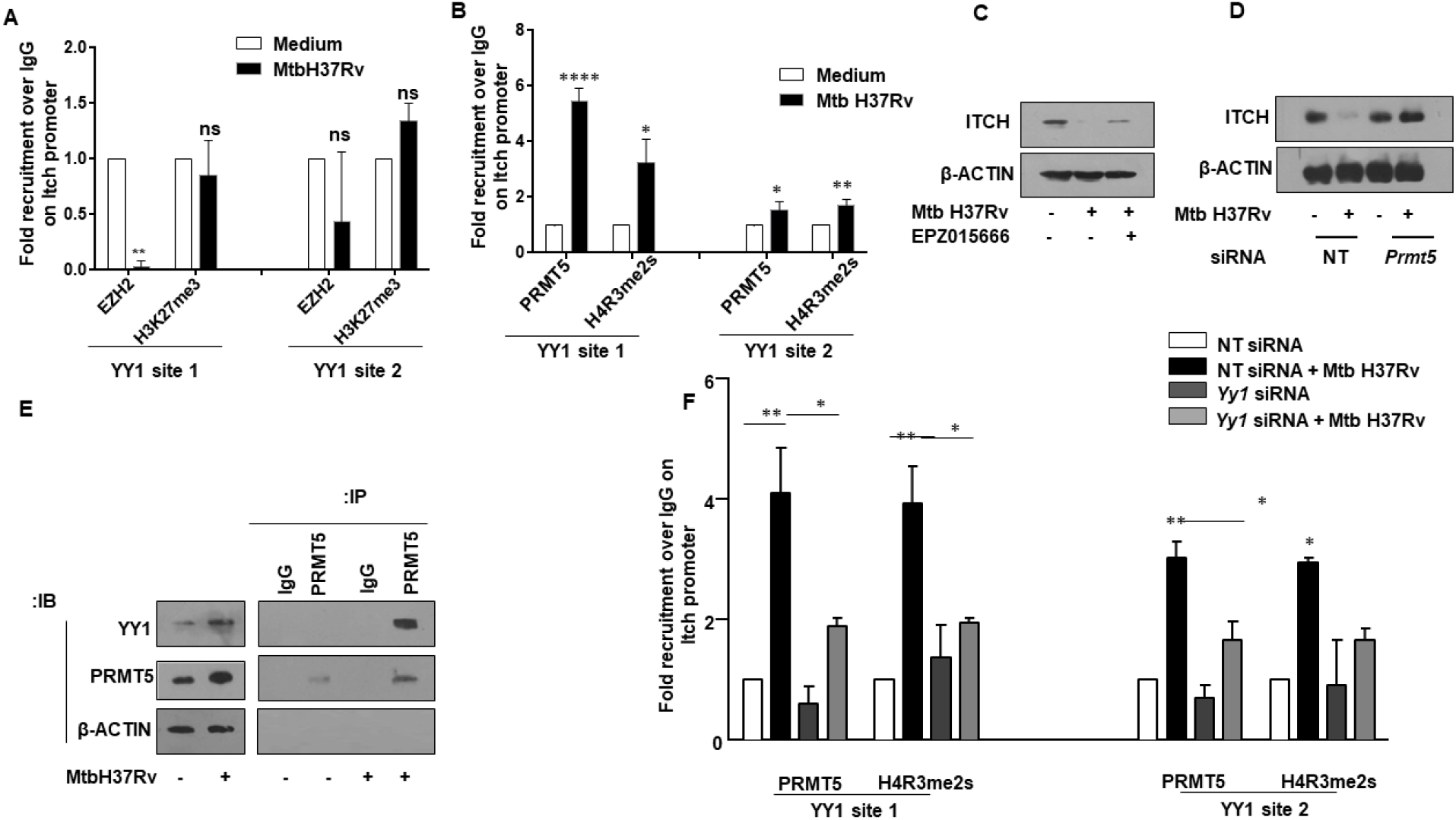
YY1 associates with the arginine methyl transferase PRMT5 to regulate repression of ITCH during Mtb infection. **(A)** Mouse peritoneal macrophages were infected with Mtb H37Rv for 24 h and assessed for the recruitment of EZH2 and associated H3K27me3 marks over the *Itch* promoter by ChIP assay. **(B)** Mouse peritoneal macrophages were treated with PRMT5 inhibitor EPZ015666 for 1 h, followed by 24 h infection with Mtb H37Rv. Whole cell lysates were assessed for the expression of ITCH by immunoblotting. **(C)** Mouse peritoneal macrophages were transfected with NT or *Prmt5* siRNAs. Transfected cells were infected with Mtb H37Rv for 24 h and assessed for the expression of ITCH by immunoblotting. **(D)** Mouse peritoneal macrophages were infected with Mtb H37Rv for 24 h, whole cell lysates were immunoprecipitated with IgG control or PRMT5 antibodies and assessed for their interaction with YY1 by immunoblotting. **(E)** Mouse peritoneal macrophages were infected with Mtb H37Rv for 24 h and assessed for the recruitment of PRMT5 and associated H4R3me2s marks over the *Itch* promoter by ChIP assay. **(F)** Mouse peritoneal macrophages were transfected with NT or *Yy1* siRNAs. Transfected cells were infected with Mtb H37Rv for 24 h and assessed for the recruitment of PRMT5 and associated H4R3me2s marks over the *Itch* promoter by ChIP assay. qRT-PCR data represents mean±S.E.M. and immunoblotting data are representative of three independent experiments. NT, non-targeting; ChIP, Chromatin immunoprecipitation. *p<0.05; **, p<0.01; *** p<0.001;****, p < 0.0001 (One way ANOVA; GraphPad Prism 6.0).

Together, these results suggest that Mtb infection mediates the repression of the E3 ligase through the concerted action of YY1 and PRMT5.

### PRMT5 activity contributes to sustained lipid accumulation in Mtb-infected macrophages

With the premise that YY1 and PRMT5 orchestrate the Mtb-mediated repression of ITCH, we assessed the role of PRMT5 in FM formation. To this end, PRMT5 expression was compromised in primary murine macrophages infected with Mtb using specific siRNA. We found that loss of PRMT5 restricted the Mtb-induced expression of ADRP and CD36, and consequently lipid accumulation **(Fig. 5A, C, D)**. In line, inhibition of PRMT5 enzymatic activity compromised the levels of CD36 and ADRP and the resultant FMs, thereby validating the essential role of PRMT5 in lipid accrual during mycobacterial infection **(Fig. 5B, E, F)**. Since accumulated lipids provide a favorable niche to the internalized mycobacteria (35), we evaluated the status of mycobacterial survival in macrophages upon treatment with PRMT5 inhibitor. Perturbation of PRMT5 enzyme activity in infected macrophages revealed a significant reduction in mycobacterial burden **(Fig. 5G)**, thereby indicating a compelling role of the PRMT5 inhibitor, EPZ015666, in reducing Mtb survival.

**Figure 5.**
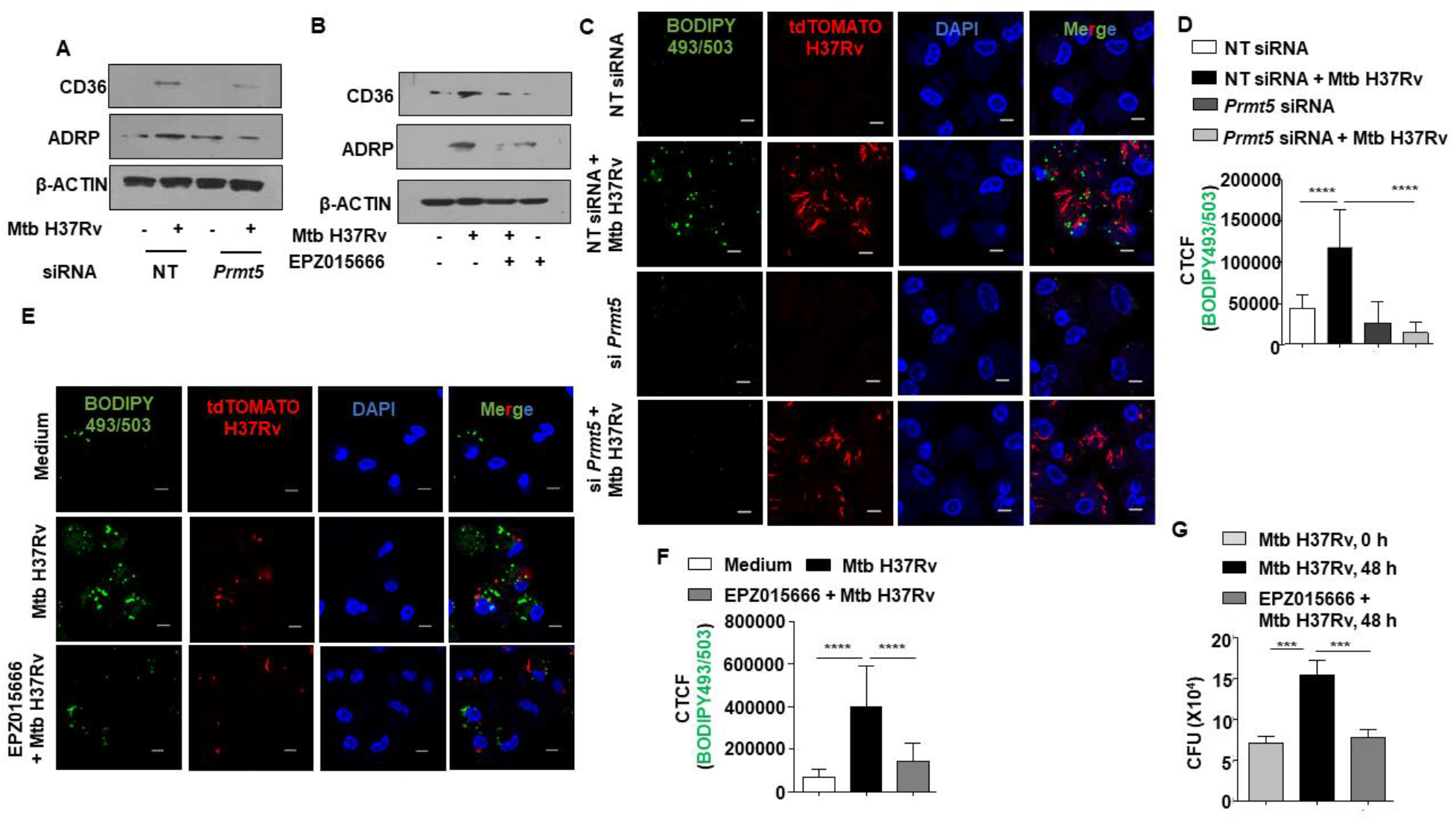
PRMT5 activity aids in the sustained accumulation of lipid droplets during Mtb infection. **(A)** Mouse peritoneal macrophages were transfected with NT or *Prmt5* siRNAs. Transfected cells were infected with Mtb H37Rv for 24 h and assessed for the expression of ADRP and CD36 by immunoblotting. **(B)** Mouse peritoneal macrophages were treated with PRMT5 inhibitor EPZ015666 for 1 h, followed by 24 h infection with Mtb H37Rv. Whole cell lysates were assessed for the expression of ADRP and CD36 by immunoblotting. **(C, D)** Mouse peritoneal macrophages were transfected with NT or *Prmt5* siRNAs. Transfected cells were infected with tdTomato Mtb H37Rv for 48 h and analyzed for lipid accumulation (BODIPY493/503) by confocal microscopy; **(C)** representative image and **(C)** respective quantification. **(E, F)** Mouse peritoneal macrophages were treated with PRMT5 inhibitor EPZ015666 for 1 h, followed by 48 h infection with tdTomato Mtb H37Rv and analyzed for lipid accumulation (BODIPY493/503) by confocal microscopy; **(E)** representative image and **(F)** respective quantification. **(G)** Mouse peritoneal macrophages were infected with Mtb H37Rv for 4 h. Extracellular bacteria were removed, and the infected cells were cultured in the presence or absence of PRMT5 inhibitor EPZ015666 for 48 h. Cells were lysed and plated on 7H11 to enumerate intracellular Mtb H37Rv burden. All immunoblotting and immunofluorescence data are representative of three independent experiments. NT, non-targeting; CTCF, corrected total cell fluorescence; CFU, colony forming units. *** p<0.001;****, p < 0.0001 (One way ANOVA; GraphPad Prism 6.0).

### Perturbation of PRMT5 activity aids in enhanced mycobacterial killing and resolution of granuloma-like lesions during infection

Having established the role of YY1-PRMT5 axis in mediating ITCH repression and consequently FM formation during Mtb infection, we employed an *in vivo* mouse model of TB **(Fig. 6A)** to understand the effect of perturbing the signaling axis on mycobacterial organ burden and pulmonary pathology. Utilizing the specific pharmacological inhibitor, EPZ015666, we found that perturbation of PRMT5 enzyme activity compromised FM generation in the lungs of infected mice **(Fig. 6B)**. Further, the formation of hallmark TB granulomatous lesions within the lungs of treated mice was significantly reduced as indicated by the H and E-stained lung sections and assessment of granuloma fraction within the infected lungs **(Figure 6C, D)**. Corroborating our histological findings, we observed an appreciable decrease in mycobacterial burden in the lungs of Mtb-infected mice treated with PRMT5 inhibitor.

**Figure 6.**
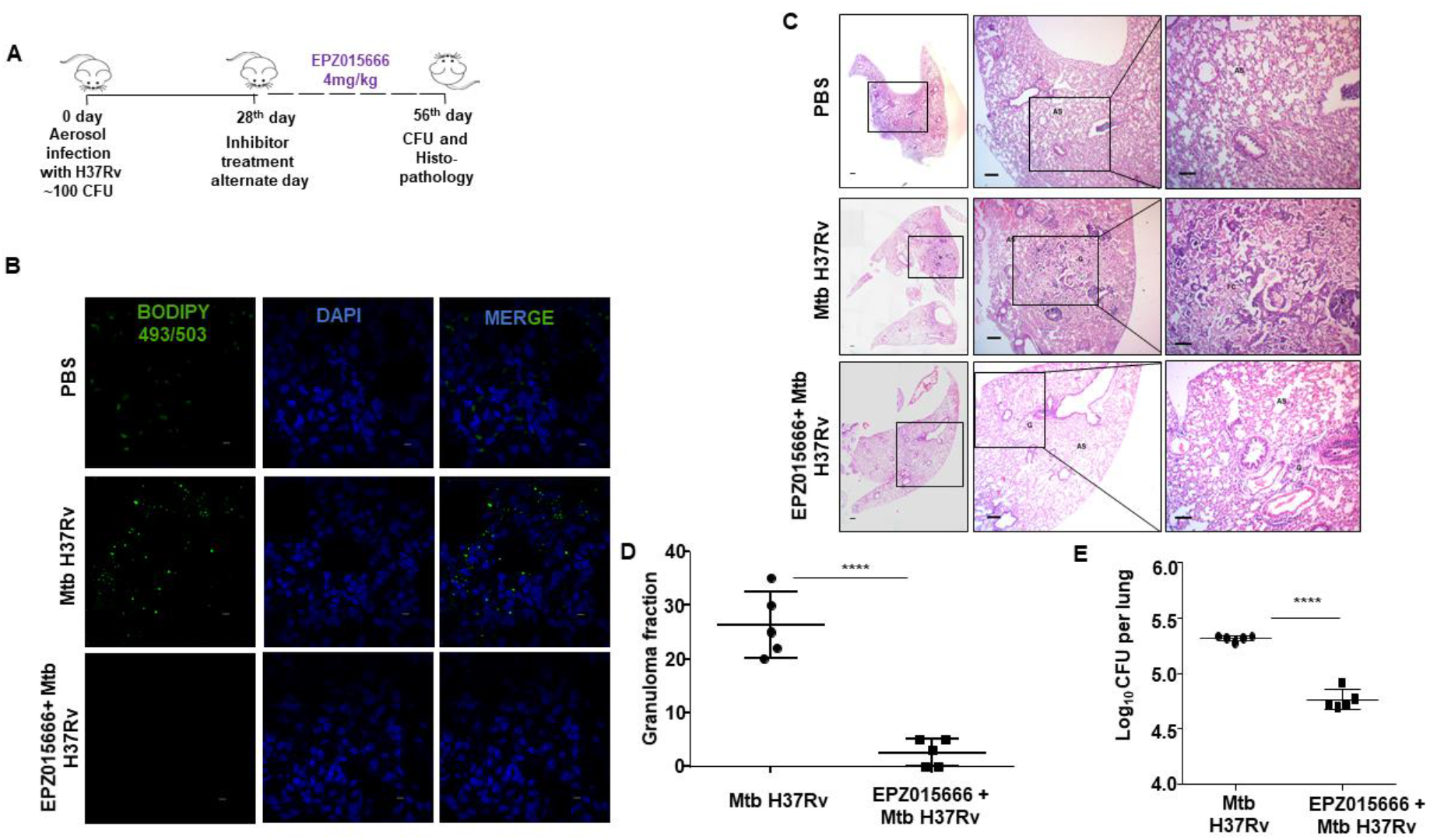
Inhibition of PRMT5 alleviates TB pathology. **(A)** Schematic representing the therapeutic regimen followed in a mouse model of Mtb infection. **(B-E)** Balb/c mice were infected with Mtb H37Rv and treated with PRMT5 inhibitor EPZ015666 as indicated in (A). **(B)** Lung cryosections were assessed for lipid accumulation by confocal microscopy (BODIPY493/503) (number of mice per group = 3). **(C, D)** TB pathology (granulomatous lesions) in the lungs was analyzed by H and E staining; **(C)** representative images and **(D)** respective quantification (number of mice per group = 5). **(E)** Mycobacterial burden in the lungs of infected and PRMT5 inhibitor treated mice was enumerated by plating lung homogenates on 7H11 plates (number of mice per group = 5). Specific regions of the H and E-stained sections were zoomed by the pathologist for evaluation of granuloma fraction. Accordingly, the portions that have been zoomed are demarcated in the images; Magnification, 4X (left panel); 40X (middle panel); 100X (right panel); Scale bar for H and E images, 200 μm. Scale bar of immunofluorescence, 5μm. G, granulomatous lesion; AS, alveolar space, FC, foam cell; CFU, colony forming units. ****, p < 0.0001 (Student’s t-test; GraphPad Prism 6.0).

Thus, we first report the YY1-PRMT5-dependent repression of the E3 ligase, ITCH, during Mtb infection. The reduction in the expression of ITCH contributed to enhanced lipid accretion in infected cells, thereby aiding mycobacterial survival, both *in vitro* and *in vivo.* With these lines of evidence, we highlight the crucial contribution of NOTCH-YY1-PRMT5 axis in the pathogenesis of TB disease **(Fig. 7)**.

**Figure 7.**
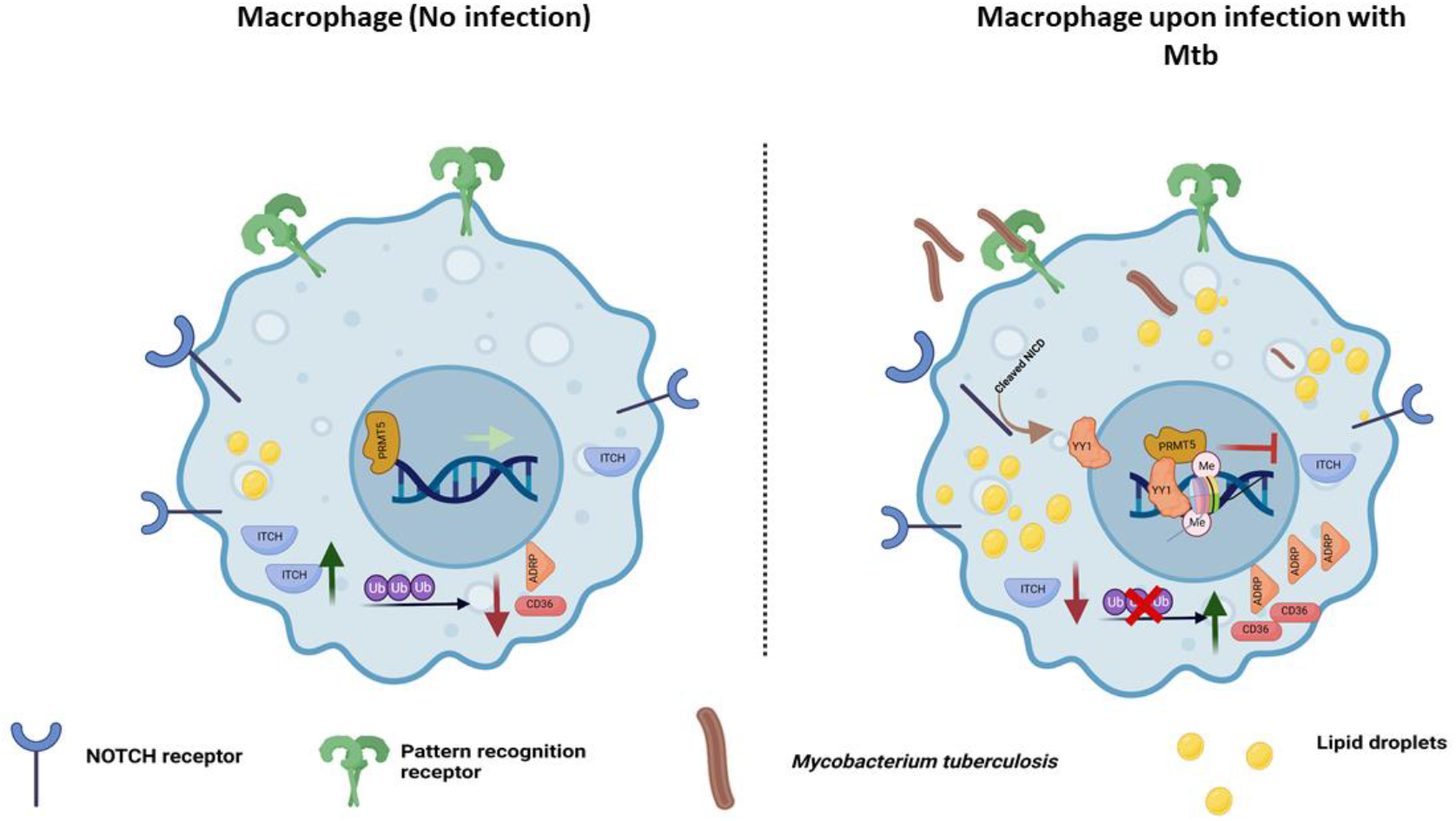
Model: Mycobacteria utilizes YY1-PRMT5 axis to repress ITCH that aids in lipid accumulation.

## Discussion

The enormous success of Mtb in continuing to affect human lives stems from its ability to persist within the body of the infected individual and attain adequate insurance against definitive drugs. FM formation has been attributed as a major development that aids pathogen survival within the hypoxic environment of the host cell. Untreated TB patients harbor lipid rich FMs in their lungs that have been shown to contain viable mycobacteria (36). Mtb-induced FMs are rich in triacylglycerols and sterol esters which become the major source of nutrition for the pathogen wherein it slowly acquires a dormant phenotype (37). While metabolic adjustments by the bacterium in acquiring a dormancy phenotype provides survival advantages, FMs provide additional benefits to Mtb for its sustenance. Interestingly, FMs displayed enhanced protection from cell death (38) whilst also demonstrating a change to a more favorable inflammatory milieu for mycobacterial survival (4, 38). Since several proteins have been implicated in Mtb-driven FM formation, it becomes pertinent to explore the regulatory mechanisms that contribute to the expression and stability of these proteins within the cells.

Ubiquitination is implicated in diverse cellular processes including protein degradation by the proteasome, cell cycle progression, transcriptional regulation, DNA repair and signal transduction (39). Pathogenic species have devised crafty ways to hijack this host cellular process to augment their own survival. Distinct bacteria such as *Salmonella, Shigella* and *Legionella* secrete ubiquitin ligase-like effectors which are involved in the modulation of critical events of the host that subsequently aid in pathogen survival (40). Specifically, *Salmonella enterica* could effectuate ubiquitination of MHCII through its effector protein SteD, which acts as an adaptor between MHCII and the host E3 ligase MARCH8 (41). Also, *Legionella pneumophila* has been shown to mediate the proteasomal degradation of several proteins assembled on the Legionella containing vacuole (LCV) through its effector protein AnkB. Besides, changes in the expression of distinct host E3 ligases could contribute to bacterial pathogenesis. Mice deficient in the E3 ligase gene *Hectd3* could enhance host immune responses against bacteria such as *Francisella novicida, Mycobacterium bovis* (BCG), and *Listeria* and limit their dissemination (42). Furthermore, as mentioned earlier, NEDD4 (a member of the HECT family of E3 ligases) contributed to enhanced autophagy and mitophagy in infected cells, thereby leading to bacterial clearance (14). In the current study, analysis of the expression of the NEDD4-family of E3 ligases revealed an appreciable decline in the expression of ITCH. While we established the role of ITCH in contributing to pro-mycobacterial FM formation, its role in modulating anti-mycobacterial processes require further investigation. The role of EGFR pathway in augmenting Mtb survival has been underscored previously (6, 43). Available reports allude towards a direct effect of ITCH on the stability of EGF tyrosine kinase receptor, thereby bringing about EGFR signaling downregulation (44). Thus, repression of ITCH could be yet another mechanism that contributes towards EGFR signaling activation during Mtb infection. Besides, ITCH has also been shown to regulate the turnover of distinct proteins involved in coordinating T cell immunity (45). Altogether, additional consequences of Mtb-mediated downregulation of ITCH at the systemic scale other than its role in FM generation, would be promising avenues for future research.

Mtb infection launches a circumspect regulation of the host cell transcriptional regulators to effectuate lipid accumulation in macrophages. Congruent to earlier reports (28), the expression of YY1 was found to be elevated in the current study upon virulent mycobacterial infection. In parallel studies, targeted knockdown of YY1 decreased the expression of several key effectors of lipid metabolism (46). Besides, YY1 promoted lipid accumulation in zebrafish liver (47). Additionally, YY1 has been reported to play a significant role in different infectious scenarios. Notably, IFN-1 production is under the dynamic regulation of YY1 during different stages of viral infection (48). YY1 could enhance the gene expression of Human T lymphotropic virus type 1 (HTLV-1) by binding to specific viral RNA (49). Furthermore, YY1 aided in the integration of Moloney Murine Leukemia Virus cDNA into the host chromosomes (50). Thus, it was of our interest to evaluate the role of YY1 in regulating pro-mycobacterial events in infected cells. Moreover, we also attempted to dissect the definitive signaling events that led to the enhanced expression of YY1. Albeit our initial observations designated a role for the NOTCH signaling pathway in the regulation of YY1, available literature also indicated the possible association between NOTCH pathway and YY1 upregulation. To our interest, CBF-1 independent NOTCH signaling could modulate the gene expression of YY1 target genes (51). An ancillary support for the role of NOTCH pathway could be derived from two reports wherein the juxtacrine signaling pathway was instrumental in *M. bovis-* mediated SOCS3 upregulation in macrophages on one hand (52), while on the other, YY1 was important for the elevated expression of SOCS3 in neuroinflammation and neuropathic pain (53).

During infection, the surface proximal events of signaling activation are transformed into distinct immunological responses via the action of cellular intermediates that regulate nuclear transactions, including transcription factors and epigenetic modulators. Regulated histone methylation has a diverse role to play during several viral and bacterial infections. For instance, influenza virus and vesicular stomatitis virus have been shown to induce H3K79 methylation by DOT1 histone methyl transferase (HMT) that controls antiviral interferon signaling in host cells (54). The HMT SET8 has been reported to modulate apoptosis and inflammatory responses during mycobacterial infection (18), while EZH2 has been implicated in the downmodulation of antigen presentation (55). Further, demethylases such as JMJD3 and acetylation readers such as BRD4 have been shown to regulate pathogen-specific lipid accumulation (4, 6). Here, we uncover the role of PRMT5 in mediating lipid accumulation through its close association with the transcriptional regulator, YY1. Histone methyl transferases, including those belonging to the arginine methyl transferase family (PRMTs) have been shown to associate with YY1. Precisely, PRMT1 and PRMT7 interact with YY1 to regulate the expression of specific genes (33, 56). Besides, YY1 could transcriptionally activate PRMT5 and aid in proliferation and invasion in laryngeal cancer cells (57). With our observation on the close association between PRMT5 and YY1, it would be worthwhile in future to understand the co-occupancy of the two factors on distinct gene loci that contribute to mycobacterial survival within macrophages.

Taken together, the current study unravels the implication of the interplay of specific signaling components and host epigenetic machinery in governing the pathogenesis of Mtb. The bacterium is highly intricate in its mechanisms of infection and presents itself in drug-resistant forms, thereby posing definite bottlenecks in the development of effective therapeutics. In this light, it becomes highly important to study the exact molecular details of infection and understand the roles that the host play in driving its pathogenesis to create a library of possible target molecules for therapeutic purposes.

Although the regulation of lipid accumulation has been studied at the transcriptional level, we provide an alternate perspective of the relevance of turnover of lipids. We show that controlling the turnover of proteins associated with FMs via proteasomal degradation is important to aggregate lipids and that the downregulation of the E3 ubiquitin ligase ITCH by YY1-PRMT5 axis plays a primary role in this process. Besides, the lipids can also undergo turnover by distinct regulated processes such as enzymatic degradation (by lipases) and lipid-specific autophagy. Uncovering the contribution of each of these mechanisms would be useful in designing the most effective combination of drug targets against Mtb infection. Additionally, the effect of the use of the PRMT5 inhibitor in reducing Mtb burden requires further investigation. PRMT5 has been reported to negatively regulate cGAS-mediated antiviral responses (58). It must be noted that available reports have alluded to the role of PRMT5 in regulating the alternative splicing of genes (59). In this respect, Mtb infection has been shown to regulate alternate splicing events to program macrophage responses (60, 61), The reduced mycobacterial burden and granuloma-like regions within the lungs of infected mice upon PRMT5 inhibitor administration might be indicative of a differential immune response to the pathogen when compared to the untreated animals. While reduced lipids would certainly contribute to decreased mycobacterial burden, it would be imperative to evaluate the role of PRMT5 inhibitor in deregulating the alternative splicing events and modulating the other discussed immune mechanisms during mycobacterial infection.

## Materials and Methods

### Cells and Mice

Four to six weeks old, male and female BALB/c mice were utilized for all experiments. Mice were procured from The Jacksons Laboratory and maintained at the Central Animal Facility (CAF) in Indian Institute of Science (IISc) under 12-hour light and dark cycle. For *in vitro* experiments, mouse peritoneal macrophages were utilized. Briefly, mice were injected intraperitoneally with (4-8 %) Brewer’s thioglycollate and peritoneal exudates were harvested in ice cold PBS after four days, seeded in tissue culture dishes. Adherent cells were utilized as peritoneal macrophages. RAW 264.7 mouse monocyte-like cell line was obtained from National Centre for Cell Sciences (NCCS), India. All cells were cultured in Dulbecco’s Modified Eagle Medium (DMEM, Gibco, Thermo Fisher Scientific) supplemented with 10% heat inactivated Fetal Bovine Serum (FBS, Gibco, Thermo Fisher Scientific) and maintained at 37 °C in 5% CO_2_ incubator.

### Ethics Statement

Experiments involving mice were carried out after the approval from Institutional Ethics Committee for animal experimentation. The animal care and use protocol adhered were approved by national guidelines of the Committee for the Purpose of Control and Supervision of Experiments on Animals (CPCSEA), Government of India. Experiments with virulent mycobacteria (Mtb H37Rv) were approved by the Institutional Biosafety Committee.

### Bacteria

Virulent strain of Mtb (Mtb H37Rv) was a kind research gift from Prof. Amit Singh, Department of Microbiology and Cell Biology, and Centre for Infectious Disease Research, IISc. Mycobacteria were cultured in Middlebrook 7H9 medium (Difco, USA) supplemented with 10% OADC (oleic acid, albumin, dextrose, catalase). Single-cell suspensions of mycobacteria were obtained by passing mid log phase culture through 23-, 28- and 30-gauge needle 10 times each and used for infecting primary cells or RAW 264.7 cells at multiplicity of infection 10. The studies involving virulent mycobacterial strains were carried out at the biosafety level 3 (BSL-3) facility at Centre for Infectious Disease Research (CIDR), IISc.

### Reagents and antibodies

All general chemicals and reagents were procured from Sigma-Aldrich/ Merck Millipore, HiMedia and Promega. Tissue culture plastic ware was purchased from Jet Biofil or Tarsons India Pvt. Ltd. and Corning Inc. siRNAs were obtained from Dharmacon as siGENOME SMART-pool reagents against *Yy1*, *Prmt5, Notch1.* Oleic acid, HRP-tagged anti-β-ACTIN (A3854), 4’,6-Diamidino-2-phenylindole dihydrochloride (DAPI) were procured from Sigma-Aldrich. Anti-cleaved NOTCH1, anti-ITCH, anti-YY1, anti-α-TUBULIN, anti-PRMT5, anti-EZH2, anti-H3K27me3 antibodies were procured from Cell Signaling Technology (USA). Anti-H4R3me2s antibody was sourced from Abcam. Anti-ADRP and anti-CD36 antibodies were procured from Santa Cruz Biotechnology (USA). Anti-LAMINB antibody was purchased from IMGENEX. HRP conjugated anti-rabbit IgG/ anti-mouse IgG was obtained from Jackson ImmunoResearch (USA). Lipofectamine 3000 was purchased from Thermo Fisher Scientific. BODIPY 493/503 (4,4-Difluoro-1,3,5,7,8-Pentamethyl-4-Bora-3a,4a-Diaza-s-Indacene lipid stain was from Molecular Probes (Invitrogen/ Thermo Fisher Scientific).

### Treatment with pharmacological reagents

Primary macrophages pre-treated with the following reagents one hour prior to infection with Mtb H37Rv: GSI (Calbiochem, 10 μM); EPZ015666 (Tocris, 20 μM); Gefitinib (Cayman Chemicals, 20 μM); LY294002 (Calbiochem, 50 μM); IWP-2 (Calbiochem, 20 μM). MG132 (Sigma-Aldrich, 10 μM) was added 4 h. prior to harvest.

### Plasmids and constructs

NICD overexpression (OE) plasmid and β-galactosidase plasmids were received as kind gifts from Prof. Kumaralvel Somasundaram, (IISc, Bangalore). ITCH OE plasmid was procured from Addgene. HES1-luciferase plasmid was a kind gift from Prof. Ryoichiro Kageyama, Institute for Virus Research, Kyoto University.

### Transient transfection studies

RAW 264.7 macrophages were transfected with the indicated constructs (NICD OE, ITCH OE, HES1-luc.); or primary macrophages were transfected with 100 nM each of siGLO Lamin A/C, non-targeting siRNA or specific siRNAs with the help of Lipofectamine 3000 for 6 h; followed by 24 h recovery. 70-80 % transfection efficiency was observed by counting the number of siGLO Lamin A/C positive cells in a microscopic field using fluorescence microscopy. Transfected cells were subjected to the required infections/ treatments for the indicated time points and processed for analyses.

### Luciferase Assay

RAW 264.7 cells were transfected with HES1-luciferase and β-galactosidase plasmids using Lipofectamine 3000 for 6 h, followed by 24 h of recovery. subjected to the required infections/ treatments for the indicated time points and processed for analyses. Briefly, cells were harvested and lysed in reporter lysis buffer (Promega) and luciferase activity was assayed using luciferase assay reagent (Promega). The results were normalized for transfection efficiencies by assay of β-galactosidase activity

### RNA isolation and quantitative real time PCR (qRT-PCR)

Treated samples were harvested in TRIzol (Sigma-Aldrich) and incubated with chloroform for phase separation. Total RNA was precipitated from the aqueous layer. Equal amount of RNA was converted into cDNA using First Strand cDNA synthesis kit (Applied Biological Materials Inc.). The cDNA thus obtained was used for SYBR Green (Thermo Fisher Scientific) based quantitative real time PCR analysis for the concerned genes. *Gapdh* was used as internal control gene. Primer pairs used for expression analyses are provided below:

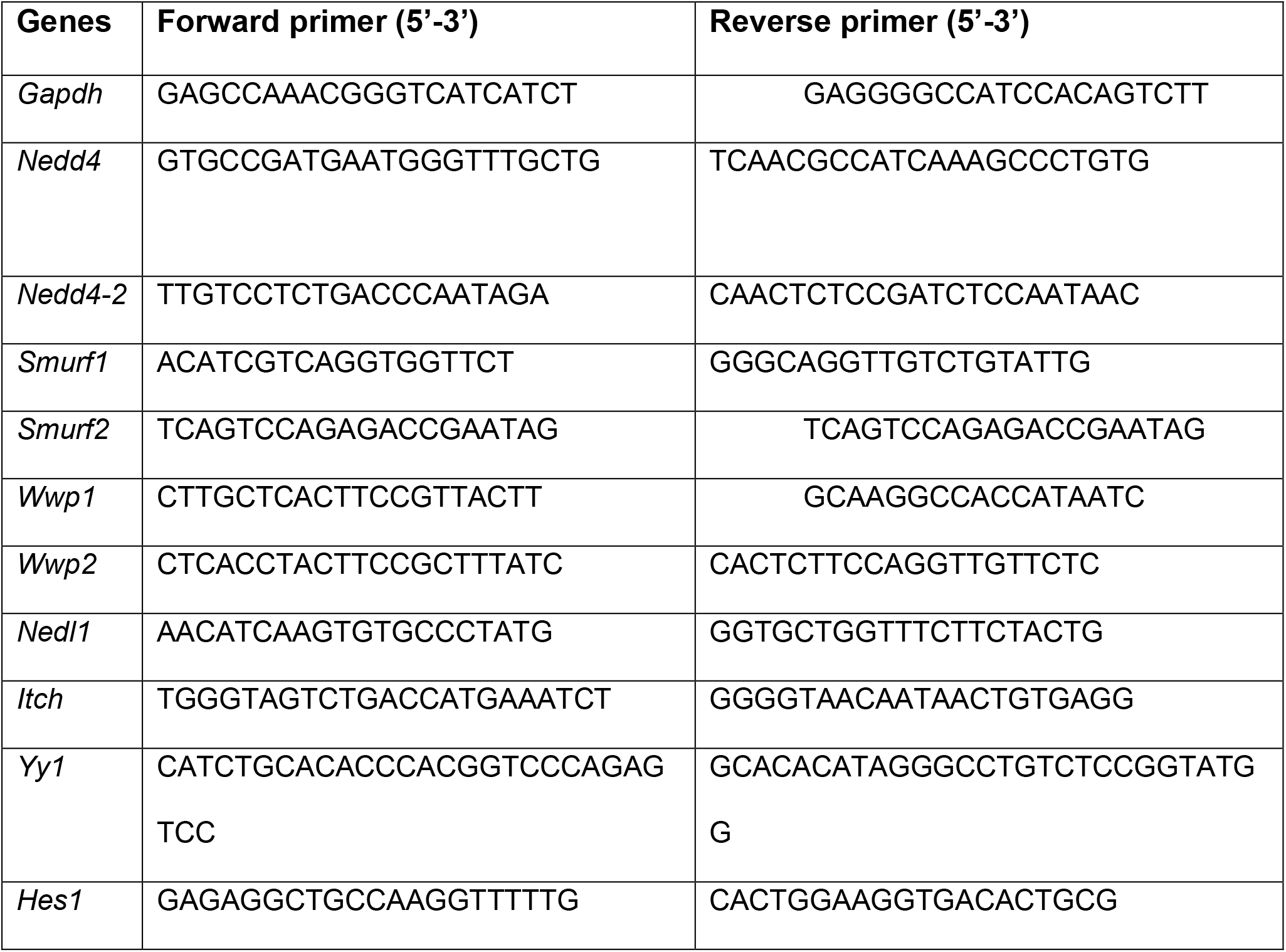

Primers were synthesized and obtained from Eurofins Genomics Pvt. Ltd. (India).

### Nuclear-cytoplasmic extraction

Cells were treated as indicated, harvested by centrifugation, and gently resuspended in ice-cold Buffer A (10 mM HEPES pH 7.9, 10 mM KCl, 0.1 mM EDTA, 0.1 mM EGTA, 1 mM DTT, and 0.5 mM PMSF). After incubation on ice for 15 min, cell membranes were disrupted with 10% NP-40 and the nuclear pellets were recovered by centrifugation at 13,226 × g for 15 min at 4°C. The supernatants from this step were used as cytosolic extracts. Nuclear pellets were lysed with ice-cold Buffer C (20 mM HEPES pH7.9, 0.4 M NaCl, 1 mM EDTA, 1 mM EGTA, 1 mM DTT, and 1 mM PMSF) and nuclear extracts were collected after centrifugation at 13,226 × g for 20 min at 4°C.

### Immunoblotting

Cells post treatment and/or infection were washed with 1X PBS. Whole cell lysate was prepared by lysing in RIPA buffer [50 mM Tris-HCl (pH 7.4), 1% NP-40, 0.25% sodium deoxycholate, 150 mM NaCl, 1 mM EDTA, 1 mM PMSF, 1 μg/ml each of aprotinin, leupeptin, pepstatin, 1 mM Na3VO4, 1 mM NaF] on ice for 30min. Total protein from whole cell lysates was estimated by Bradford reagent. Equal amount of protein was resolved on 12% SDS-PAGE and transferred onto PVDF membranes (Millipore) by semi-dry immunoblotting method (Bio-Rad). 5% non-fat dry milk powder in TBST [20 mM Tris-HCl (pH 7.4), 137 mM NaCl, and 0.1% Tween 20] was used for blocking nonspecific binding for 60 min. After washing with TBST, the blots were incubated overnight at 4°C with primary antibody diluted in TBST with 5% BSA. After washing with TBST, blots were incubated with anti-rabbit IgG secondary antibody conjugated to HRP antibody (111-035-045, Jackson ImmunoResearch) for 4h at 4°C. The immunoblots were developed with enhanced chemiluminescence detection system (Perkin Elmer) as per manufacturer’s instructions. For developing more than one protein at a particular molecular weight range, the blots were stripped off the first antibody at 60 °C for 5 min using stripping buffer (62.5 mM Tris-HCl, with 2 % SDS 100 mM 2-Mercaptoethanol), washed with 1X TBST, blocked; followed by probing with the subsequent antibody following the described procedure. β-ACTIN was used as loading control.

### Immunoprecipitation Assay

Immunoprecipitation assays were carried out following a modified version of the protocol provided by Millipore, USA. Treated samples were washed in ice cold PBS and gently lysed in RIPA buffer. The cell lysates obtained were subjected to pre-clearing with BSA-blocked Protein A beads (Bangalore Genei, India) for 30 min at 4 °C and slow rotation. The amount of protein in the supernatant was quantified and equal amount of protein was used for pull down from each treatment condition; using Protein A beads pre-conjugated with the antibody of interest or isotype control IgG antibody. After incubation of the whole cell lysates with the antibody-complexed beads for 4 h at 4 °C on slow rotation, the pellet containing the bead-bound immune complexes were washed with RIPA buffer twice. The complexes were eluted by boiling the beads in Laemmli buffer for 10 min. The bead free samples were resolved by SDS-PAGE and the target interacting partners were identified by immunoblotting. Clean-Blot™ IP Detection Reagent (21230) was obtained from Thermo Scientific.

### Chromatin Immunoprecipitation (ChIP) Assay

ChIP assays were carried out using a protocol provided by Upstate Biotechnology and Sigma-Aldrich with certain modifications. Briefly, treated samples were washed with ice cold 1X PBS and fixed with 3.6 % formaldehyde for 15 min at room temperature followed by inactivation of formaldehyde with 125 mM glycine. Nuclei were lysed in 0.1% SDS lysis buffer [50 mM Tris-HCl (pH 8.0), 200 mM NaCl, 10 mM HEPES (pH 6.5), 0.1 % SDS, 10 mM EDTA, 0.5 mM EGTA, 1 mM PMSF, 1 μg/ml of each aprotinin, leupeptin, pepstatin, 1 mM Na3VO4 and 1 mM NaF]. Chromatin was sheared using Bioruptor Plus (Diagenode, Belgium) at high power for 70 rounds of 30 sec pulse ON and 45 sec pulse OFF.

Chromatin extracts containing DNA fragments with an average size of 500 bp were immunoprecipitated with KLF5 or BRD4 or H3K27Ac or rabbit preimmune sera complexed with Protein A agarose beads (Bangalore Genei). Immunoprecipitated complexes were sequentially washed with Wash Buffer A, B and TE [Wash Buffer A: 50 mM Tris-HCl (pH 8.0), 500 mM NaCl, 1 mM EDTA, 1 % Triton X-100, 0.1 % Sodium deoxycholate, 0.1 % SDS and protease/phosphatase inhibitors; Wash Buffer B: 50 mM Tris-HCl (pH 8.0), 1 mM EDTA, 250 mM LiCl, 0.5 % NP-40, 0.5 % Sodium deoxycholate and protease/phosphatase inhibitors; TE: 10 mM Tris-HCl (pH 8.0), 1 mM EDTA] and eluted in elution buffer [1 % SDS, 0.1 M NaHCO3]. After treating the eluted samples with RNase A and Proteinase K, DNA was purified and precipitated using phenol-chloroform-ethanol method. Purified DNA was analyzed by quantitative real time RT-PCR. All values in the test samples were normalized to amplification of the specific gene in Input and IgG pull down and represented as fold change in modification or enrichment. All ChIP experiments were repeated at least three times. The list of primers is given below:

Primers were synthesized and obtained from Eurofins Genomics Pvt. Ltd. (India).

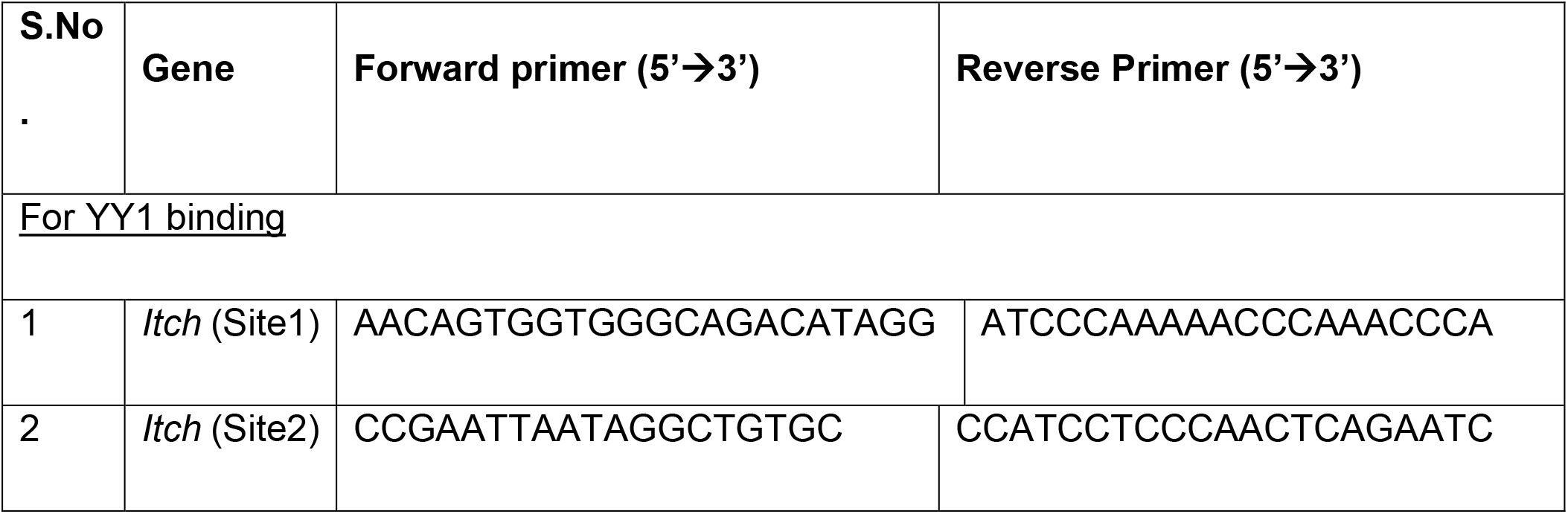

### *In vitro* CFU analysis

Mouse peritoneal macrophages were infected with Mtb H37Rv at MOI 5 for 4 h. Post 4 h, the cells were thoroughly washed with PBS to remove any surface adhered bacteria and medium containing amikacin (0.2 mg/ml) was added for 2 h to deplete any extracellular mycobacteria. After amikacin treatment the cells thoroughly washed with PBS were taken for 0 h time point, and a duplicate set was maintained in antibiotic free medium for next 48 h along with respective inhibitors GSI and EPZ015666. Intracellular mycobacterial burden was enumerated by lysing macrophages with 0.06 % SDS in 7H9 Middlebrook medium. Appropriate dilutions were plated on Middlebrook 7H11 agar plates supplemented with OADC (oleic acid, albumin, dextrose, catalase). Total colony forming units (CFUs) were counted after 21 days of plating.

### *In vivo* mouse model for TB and treatment with pharmacological inhibitor

BALB/c mice (n=30) were infected with mid-log phase Mtb H37Rv, using a Madison chamber aerosol generation instrument calibrated to 100 CFU/animal. Aerosolized animals were maintained in securely commissioned BSL3 facility. Post 28 days of established infection, mice were administered eight intra-peritoneal doses (modified from (62)) of EPZ015666 (4mg/kg) every alternate day over 28 days. On 56^th^ day post inhibitor treatment, mice were sacrificed, the left lung lobe was homogenized in sterile PBS, serially diluted, and plated on 7H11 agar containing OADC to quantify CFU. Upper right lung lobes were fixed in formalin, and processed for hematoxylin and eosin staining, or immunofluorescence analyses. Also, specific lobes from the lungs of mice were homogenized for the extraction of RNA and protein.

### Hematoxylin and Eosin staining

Microtome sections (5 μm) were obtained from formalin-fixed, paraffin-embedded mouse lung tissue samples using Leica RM2245 microtome. Deparaffinized and rehydrated sections were subjected to Hematoxylin staining followed by Eosin staining as per manufacturer instructions. After dehydrating, sections were mounted using permount. Sections were kept for drying overnight and handed over to consultant pathologist for blinded analyses

### Cryosection preparation

The excised lung tissue portions were fixed in 4% paraformaldehyde solution. Subsequently, the tissues were kept in 30% sucrose solution. The fixed lung pieces were placed in the optimal cutting temperature (OCT) media (Jung, Leica). Cryosections of 10 μm were prepared using Leica CM 1510 S or Leica CM 3050 S cryostat and then stored at −80 °C.

### Immunofluorescence

Cells were fixed with 3.6 % formaldehyde for 30 min. at room temperature. Fixed samples were blocked with 2 % BSA in PBST (containing 0.02% saponin) for 1 h. After blocking, samples were stained with the indicated antibodies at 4 °C overnight, followed by incubation with DyLight 488-, Alexa 549- or Alexa 643-conjugated secondary antibodies for 2 h and nuclei were stained with DAPI. The samples were mounted on glycerol. Confocal images were taken with Zeiss LSM 710 Meta confocal laser scanning microscope (Carl Zeiss AG, Germany) using a plan-Apochromat 63X/1.4 Oil DIC objective (Carl Zeiss AG, Germany) and images were analyzed using ZEN 2009 software.

### Lipid droplet staining

Lipid droplets were stained using neutral lipid dye BODIPY 493/503 (Invitrogen). Formaldehyde-fixed cells were stained with BODIPY (10 μg/ml) for 30 min at room temperature. Cells were washed, and nuclei were stained with DAPI. After washing with PBS, samples were mounted on glycerol and visualized and analyzed by confocal microscopy.

### Statistical analysis

Levels of significance for comparison between samples were determined by the student’s t-test and one-way ANOVA followed by Tukey’s multiple-comparisons. The data in the graphs are expressed as the mean ± S.E. for the values from at least 3 or more independent experiments and P values < 0.05 were defined as significant. GraphPad Prism software (6.0 versions, GraphPad Software, USA) was used for all the statistical analyses.

## Acknowledgements

We thank CAF, IISc for maintaining and providing mice for experimentation. NICD OE and β-galactosidase plasmids were kind research gifts from Prof. Kumaravel Somasundaram, Department of Microbiology and Cell Biology, IISc. HES1-luciferase plasmid was a kind gift from Prof. Ryoichiro Kageyama, Institute for Virus Research, Kyoto University. We acknowledge the help of BSL-3 facility and staff for helping us in our *in vitro* and *in vivo* experiments with Mtb H37Rv.

## Author contributions

SMB, conceptualization, investigation, formal analysis, manuscript draft preparation, editing; RSR, investigation; KNB, conceptualization, formal analysis, supervision, manuscript review and editing.

## Supplementary Figure Legends

**Figure S1:**
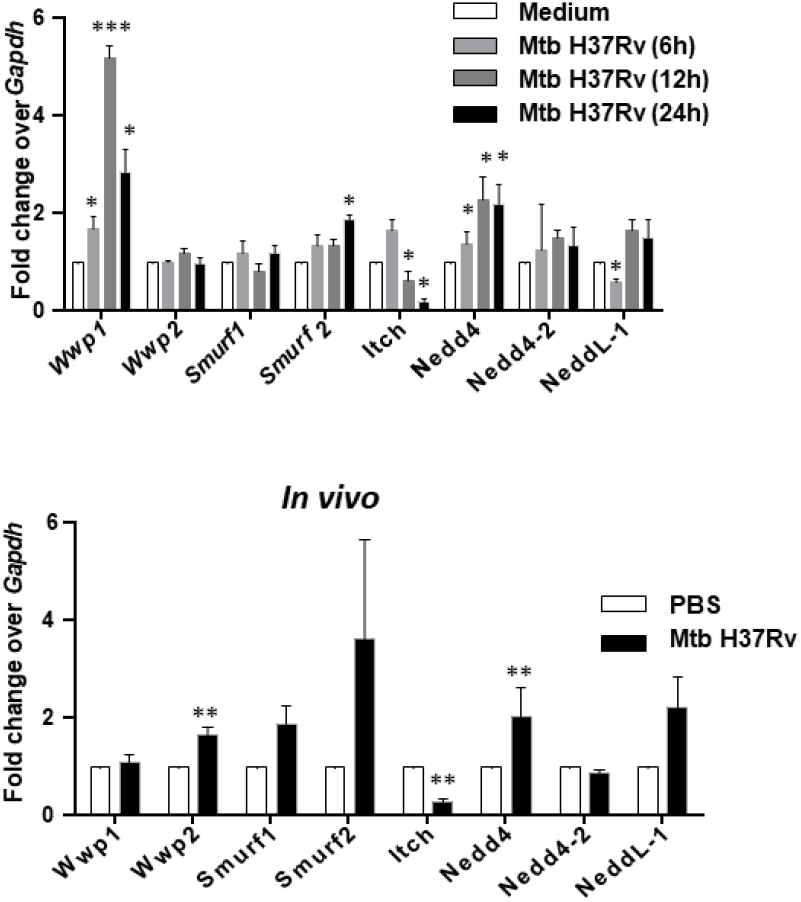
ITCH is downregulated during mycobacterial infection. **(A)** Mouse peritoneal macrophages were infected with Mtb H37Rv for the indicated time points and assessed for the transcript levels of NEDD family E3 ubiquitin ligases. **(B)** Balb/c mice were aerosol-infected with Mtb H37Rv for 28 days. Transcript levels of NEDD family E3 ubiquitin ligases was analyzed in the lung homogenates of uninfected and infected mice by qRT-PCR, (number of mice per group = 4). qRT-PCR data represents mean±S.E.M. from three independent experiments. *, p<0.05; **, p<0.01; *** p<0.001 (Student’s t-test; GraphPad Prism 6.0).

**Figure S2:**
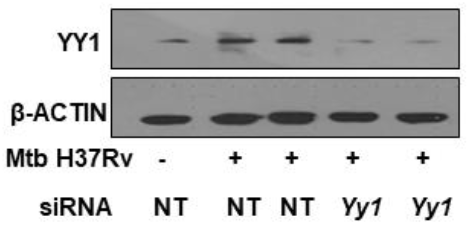
Validation of YY1 siRNA. Mouse peritoneal macrophages were transfected with NT or Yy1 siRNA. Transfected cells were infected with Mtb H37Rv for 24 h and assessed for the expression of YY1 by immunoblotting. Immunoblotting data is representative of three independent experiments. NT, non-targeting. β-ACTIN was utilized as loading control.

**Figure S3:**
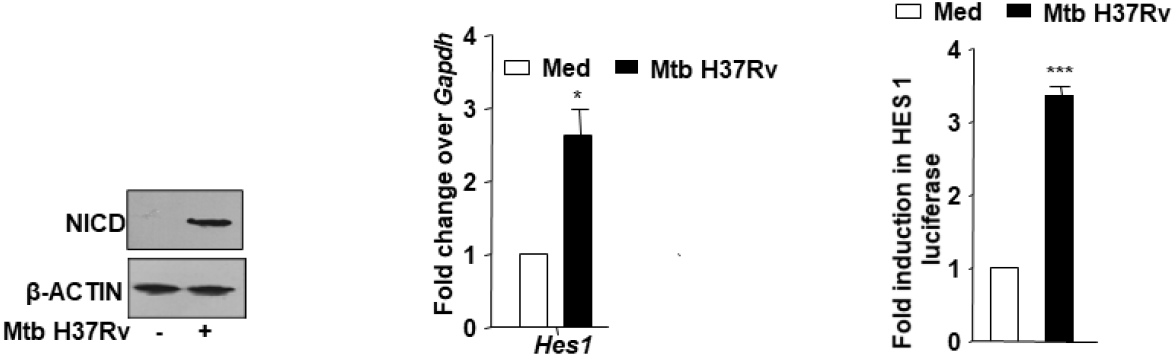
NOTCH signaling is upregulated upon mycobacterial infection. **(A)** Mouse peritoneal macrophages were infected with Mtb H37Rv for 1 h. Whole cell lysates were assessed for NICD expression by immunoblotting. **(B)** Mouse peritoneal macrophages were infected with Mtb H37Rv for 24 h and assessed for the expression of NOTCH target gene *Hes1* by qRT-PCR. **(C)** RAW264.7 macrophages were transiently transfected with HES1-luciferase and the transfected cells were infected with Mtb H37Rv for 24 h, followed by assessment of luciferase counts using luminometer. Immunoblotting data is representative of three independent experiments; qRT-PCR and luciferase data represents mean±S.E.M. NICD, NOTCH intracellular domain; Med, Medium. β-ACTIN was utilized as loading control. *, p<0.05; *** p<0.001 (Student’s t-test; GraphPad Prism 6.0).

**Figure S4:**
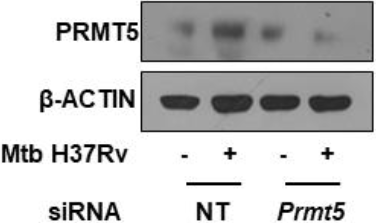
Validation of PRMT5 siRNA. Mouse peritoneal macrophages were transfected with NT or Prmt5 siRNA. Transfected cells were infected with Mtb H37Rv for 24 h and assessed for the expression of PRMT5 by immunoblotting. Immunoblotting data is representative of three independent experiments. NT, non-targeting. β-ACTIN was utilized as loading control.

